# Implicit visuomotor adaptation to clamped feedback is reduced in adults who stutter

**DOI:** 10.64898/2026.06.24.734039

**Authors:** J. Liu, K.R. Loudermilk, K.S. Kim

## Abstract

It has been demonstrated that people who stutter exhibit atypical motor control not only in speech tasks but also movements in the non-speech effector system, such as finger or arm motion. Notably, studies have reported that people who stutter show limited sensorimotor adaptation (i.e., updating subsequent movements in response to sensory errors) in both speech auditory-motor (i.e., updating speech movements in response to altered auditory feedback) and upper limb visuo-motor (i.e., updating arm movements in response to altered visual feedback) tasks. Given that speech auditory-motor adaptation is mostly if not entirely implicit (i.e., participants are unaware of the learning), it is thought that people who stutter have limited implicit adaptation in the speech effector system. It remains unclear however, whether such limited implicit learning also extends to upper limb visuomotor adaptation. Here, we examined implicit visuomotor learning in adults who stutter through the means of arm reaching adaptation to clamped visual feedback which provides a cursor that is fixed in direction (8° counterclockwise from targets) regardless of the participant’s actual hand location. All participants gradually adjusted their reach angle towards the clockwise direction, adapting in response to clamped feedback, but adults who stutter showed less adaptation compared to adults who do not stutter. In addition, computational modeling suggests that this implicit adaptation difficulties in stuttering individuals may reflect reduced error sensitivity. Together, our findings suggest that implicit sensorimotor learning difficulties in adults who stutter may generalize across multiple effector systems, providing important implications for understanding sensorimotor mechanisms underlying stuttering.

**Significance statement:** By employing the clamped visual feedback paradigm during arm reaching movements, we demonstrated that adults who stutter showed less implicit visuomotor adaptation compared to adults who do not stutter. This study provides the first evidence that implicit sensorimotor adaptation limitations in developmental stuttering generalize across multiple effector systems. Our findings not only add to a growing body of evidence that stuttering is associated with domain-general sensorimotor difficulties but also point to specific underlying processes that may lead to stuttering.

## Introduction

Although developmental stuttering is a speech fluency disorder, a growing body of evidence suggests that sensorimotor difficulties associated with stuttering may extend into various other non-speech effector systems. Numerous studies identified adults who stutter (AWS) have slower finger movement initiation and execution (e.g., Max et al., 2003; Webster, 1997), slower manual reaction times (Webster & Ryan, 1991), less bimanual coordination (e.g., W. Webster, 1988; Zelaznik et al., 1997), more variable bimanual motor timing performance (e.g., Olander et al., 2010; Toyomura et al., 2021; but also see Hilger et al., 2016 for conflicting evidence), slower and/or reduced finger sequence learning (Höbler et al., 2022; Korzeczek et al., 2020; Smits-Bandstra, De Nil, & Rochon, 2006; Smits-Bandstra, De Nil, & Saint-Cyr, 2006; Smits-Bandstra & De Nil, 2007), slower reaction time and less accurate visuomotor tracking responses (Jones et al., 2002), more disrupted hand action selection during a visuomotor working-memory task (Echeverria-Altuna et al., 2026), and inaccurate arm reaching movements (Daliri et al., 2014). Recent animal studies have also demonstrated that transgenic mice carrying stuttering-associated genetic phenotypes (*N*-acetylglucosamine-1-phosphate transferase subunits α and β, GNPTAB) exhibit non-vocal motor impairments, suggesting that stuttering-linked mutations may be disrupting motor circuits not limited to speech (Bishop et al., 2025; Millwater et al., 2025).

In our previous work, we found that sensorimotor adaptation, a type of sensorimotor learning in which movements are updated in response to perturbed sensory feedback, is limited in people who stutter (Kim et al., 2020). Interestingly, stuttering individuals showed reduced adaptation not only in a speech adaptation task, but also in an arm reaching visuomotor rotation (VMR) adaptation task, in which the visual cursor location was rotated around the center of the workspace (30° counter-clockwise relative to the hand location). Specifically, we found that AWS showed a slower initial rate of visuomotor adaptation compared to adults who do not stutter (AWNS). However, because VMR adaptation consists of multiple learning processes of distinct neural pathways (Taylor et al., 2014; Tsay et al., 2022), including an explicit component driven by task performance errors (i.e., the discrepancy between movement goal and movement outcome) as well as the implicit component driven by sensory prediction errors (Mazzoni & Krakauer, 2006, i.e., the discrepancy between predicted sensory feedback and actual sensory feedback), it remains unclear which component or components of sensorimotor adaptation (i.e., explicit vs implicit) contributed to limited adaptation in AWS.

It is possible that VMR adaptation difficulties in AWS may also indicate limitations in some implicit processes based on findings that AWS show limited *speech* sensorimotor adaptation in response to altered auditory feedback, which is an entirely implicit task (i.e., participants are unaware of the learning, Keough et al., 2013; Kim & Max, 2021; Lametti et al., 2020). In other words, speech adaptation difficulties (Daliri et al., 2018; Daliri & Max, 2018; Kim et al., 2020; Sengupta et al., 2016) reflect poor implicit learning. On the other hand, given that AWS showed a slower initial rate of arm reaching VMR adaptation, it is possible that the explicit component— which plays a dominate role in the initial phase— may be limited in stuttering individuals (McDougle et al., 2015; Tsay et al., 2022).

In this study, we begin to address this question by first examining the implicit component of visuomotor adaptation in AWS and age– and sex-matched adults who do not stutter (AWNS; *N =* 20 per group). To do so, we employed a visuomotor task-irrelevant clamped feedback paradigm (Morehead et al., 2017), in which the visual feedback of the cursor was “clamped” to a fixed direction away from the target direction, regardless of the hand location. Specifically, we manipulated the visual cursor to be fixed at an 8° counter-clockwise direction (relative to targets), regardless of hand movement direction during the clamp feedback phase, and the cursor was hidden when it approached near the target, effectively eliminating any task performance errors associated with explicit learning (Taylor et al., 2014). Additionally, a recently developed computational model of sensorimotor adaptation, Perceptual Error Adaptation (PEA) model (Z. Zhang et al., 2024), was used to fit our behavioral data to determine specific processes that contribute to limited adaptation in AWS.

## Materials and Methods

### Participants

20 AWNS (mean age *M* = 25.2 years, *SD* = 9.3 years, range = 18-53 years) and 20 AWS (mean age *M* = 25.7 years, *SD* = 8.5 years, range = 19-51 years) participated in this study. Participants were all matched in pairs by age (±3 years), sex, and handedness. In each group, there were 4 female and 16 male participants. An Edinburgh Handedness Inventory (Oldfield, 1971) assessment was performed on the day of experiment to corroborate every participant’s self-reported handedness. All participant pairs were right-handed, defined as using their right hand for the majority of surveyed tasks (> 5 of 10) with the exception of one matched pair who were both self-classified and surveyed as ambidextrous, defined as showing no consistent hand preference across Edinburgh tasks. There were 5 AWS and 3 AWNS who self-reported taking some types of medication for various diagnosed neurological, psychological, or emotional conditions. No participants had any prior history of speech-language-hearing problems beyond stuttering and were naïve to the experimental paradigm.

All participants from the stuttering group self-identified as having early onset developmental stuttering (i.e., having started stuttering before adulthood). Prior to each session, a speech sample of approximately 10 minutes was video recorded, consisting of open conversational speech and a standardized reading passage. All participants were confirmed to have stuttering by the research team on the day of experiment. After each experiment, a graduate student in a speech-language pathology program trained by a certified speech-language pathologist assessed stuttering severity for all AWS, administering the Stuttering Severity Instrument (SSI-4; Riley, 2009) for verification. Of the 20 AWS, 6 were assessed as very mild, 6 mild, 5 moderate, 2 as severe. One participant scored the SSI score of 9, which is slightly below the minimum score for the “very mild” severity (10). All participants provided written informed consent in accordance with procedures by Purdue University’s Institutional Review Board.

### Clamped feedback adaptation task

Participants were seated on a height-adjustable chair facing the KINARM End-Point (BKIN Technologies Ltd., Kingston, Canada). They were asked to hold onto the handle of the End-Point with their right hand, while resting their foreheads against the headrest of the End-Point for the entire session. In the workspace, there was a red center start position (radius 1 cm) and one of eight green targets (radius 0.75 cm) located 10 cm away from starting position at 8 different angles ranging from 0° to 315° in 45° increments (8 targets) around the center of the workspace appeared in each trial (Fig. 1A). Participants were instructed to perform rapid, center-out-and-back arm movements. Participants could not view their real hand/arm location. Instead, a white cursor displayed on a display screen represented their given hand position (Fig. 1B) and was “clamped” regardless of the hand location in some trials (Fig. 1C). Before each experiment, participants were specifically informed that the visual cursor will not match their hand location in some trials. Participants were instructed to ignore their visual cursor and to make their best effort to move their hands towards the green target every trial. To maintain rapid movements, they were repeatedly instructed to move as if they were slicing through the target throughout the experiment. To prevent task performance error-based learning, which is associated with explicit learning (Taylor et al., 2014), the cursor and target were designed to disappear once the hand moved more than 8 cm from the starting position (Fig. 1D). The cursor and target reappeared 3 seconds after reaching the target, by which time most return movements had already been completed.

**Fig. 1.**
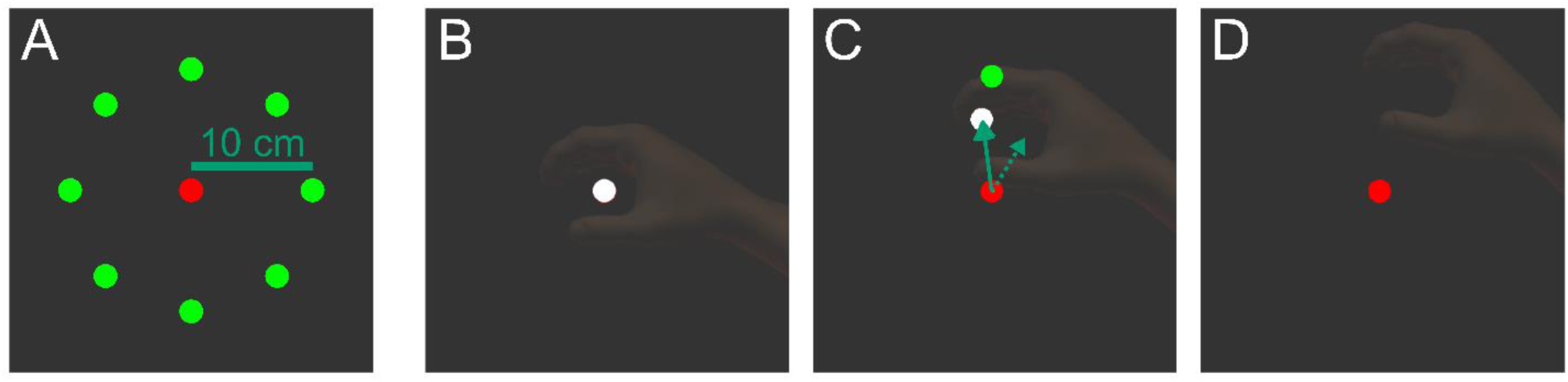
Experimental paradigm. **A**: Within every cycle, one of eight randomized green targets appeared in during which participants were instructed to reach and “slice” through the green target. **B:** The hand location was initially represented by the white cursor. **C:** During the clamped feedback trials, the white cursor location (solid arrow) was fixed to 8° counterclockwise direction from the target, regardless of the hand location (dotted arrow). **D:** Importantly, the cursor and target disappeared once the hand moved more than 8 cm from the starting point. It should be noted that participants’ view of their hand was fully occluded during the actual experiments, and the translucent hand image is shown in this figure for illustrative purposes.

There was a total of 320 trials (40 cycles, each consisting of 8 targets) in each experiment session. The first 5 cycles (cycles 1-5;40 trials) were baseline trials during which veridical feedback was presented (i.e., no perturbation), followed by 30 cycles (cycles 6-35; 240 trials) of clamped visual feedback phase during which the visual cursor maintained a fixed 8° counterclockwise direction from each target, regardless of the participant’s actual hand location. Finally, in the last five cycles (cycles 36-40; 40 trials), veridical feedback was restored. Each experiment lasted about 25 minutes.

### Data extraction and analyses

All hand position data were recorded at 1 kHz and low-pass filtered using a double-pass Butterworth filter with a 10 Hz cutoff. From the filtered position data, we calculated tangential velocity. For each trial, the movement onset was defined as the time point when the tangential velocity exceeded 5cm/s. The movement offset was defined as the time point where tangential velocity dropped below 5 cm/s or when tangential velocity reached a local minimum during the reaching movement. Peak tangential velocity was identified as the initial local maximum of the tangential velocity between the movement onset and offset.

There were a few instances during the experimental session where participants did not fully complete their initial arm reaching motion and tried to compensate by re-aiming and reaching for a second attempt during the same trial. For those few instances (about 0 to 2 trials per participant), we extracted peak velocity from their initial attempt movement (as opposed to the subsequent movement). Trials were also excluded if the tangential velocity never exceeded the 5 cm/s threshold or if a movement clearly was not aimed for the target. In total, 0.13% of trials were rejected across all participants.

We define the reach angle at each trial as the direction of a vector between start position and cursor location at the time of peak tangential velocity. The reach angle per target for all trials were normalized to baseline by subtracting the mean angle for every target across baseline cycles 3-5 (Kim et al., 2020). The amount of adaptation was then determined by the normalized reach angle (i.e., positive reach angles indicating adaptation in the clockwise direction). In this study, we examined both early and late adaptation. Early adaptation was defined as adaptation in the first 6 cycles of the clamped feedback perturbation phase (cycles 6-11, see Fig. 2A) and late adaptation as adaptation in the last 6 cycles of the perturbation phase (cycles 30-35, see Fig. 2A). For each trial that was excluded from the data, we interpolated that trial’s data point using the 2 nearest neighboring trials for that specific target (Kitchen et al., 2022). Post-processing and data organization were performed using the Signal Processing Toolbox and custom MATLAB code.

**Fig. 2.**
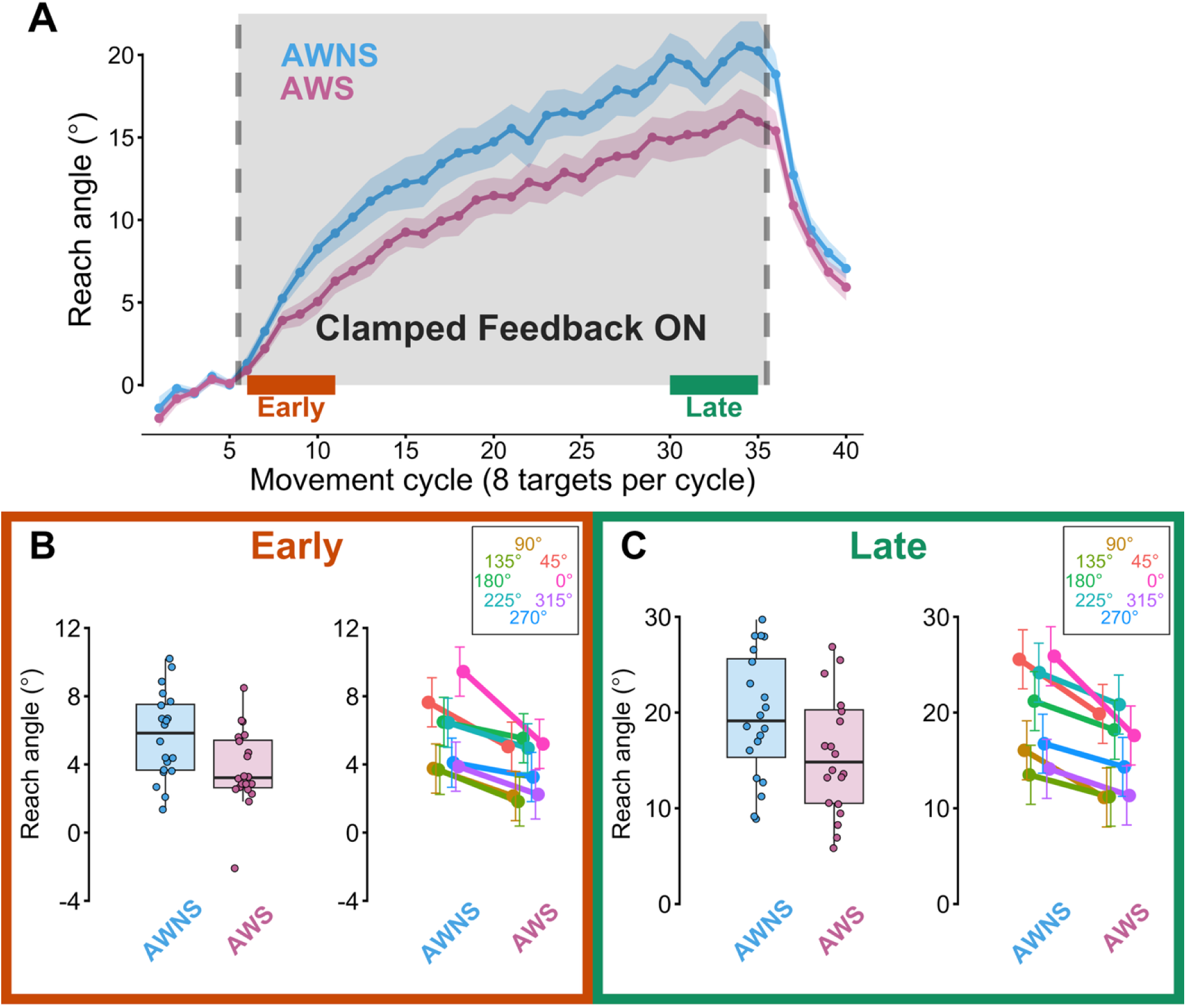
Adaptation to clamped feedback in AWS and AWNS. **(A)** Adaptation to clamped feedback in adults who stutter (AWS; pink) and adults who do not stutter (AWNS; light blue) across 40 cycles (320 trials total). Both groups’ hand reach direction changed toward clockwise direction in response to counterclockwise clamped feedback. Shaded ribbons show standard error for each group. **(B)** AWS adapted less than AWNS in early adaptation (left, cycles 6-11), p = 0.014. Late adaptation (right, cycles 30-35) also showed a significant group difference, p = 0.043. Rainbow plots represent post-hoc analyses of group × target interactions (pairwise comparisons) which revealed that AWS had reduced early adaptation in the 0° and 45° direction. In addition, AWS had reduced late adaptation when reaching for the 0°, 45°, and 90° direction. On average, AWS had a smaller extent of learning across all targets compared to AWNS, in both early and late adaptation.

### Statistical Analysis

Linear mixed-effects models using the *lmerTest* packages in R (Kuznetsova et al., 2017) were constructed. Fixed effects included group (AWS and AWNS), target (8 directions), and their interaction (group × target). The p-values and confidence intervals for the linear mixed effect model were evaluated using Satterthwaite analysis (Satterthwaite, 1941) and partial omega squared (ω^2^) values were calculated using the *omega_squared* function in the *effectsize* package (Ben-Shachar et al., 2020). For any significant interactions, post-hoc pairwise comparisons and cohen’s d effect sizes were conducted using the *emmeans* package (Lenth et al., 2026). For movement duration statistics, all between group statistical comparisons were completed with Welch’s t-tests using built in R function (R Core Team, 2025) and effect sizes were calculated using the cohen_d function in the *effectsize* package (Ben-Shachar et al., 2020).

### Computational Modeling

We utilized the PEA model for characterizing implicit adaptation. The model dynamically updates a hand angle estimate across each trial (Equation 1):

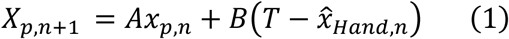

where *X_p_*_,*n*+1_ is the predicted hand angle for subsequent trials (n + 1), *A* is the retention factor, *B* is the learning rate, *T* is the target direction (fixed at 8° in this study), and *x*^*_Hand_*_,*n*_is the perceived estimated hand location that is driven by three noise sensory cues (sensory prediction, proprioception, and visual, see Equation 2):

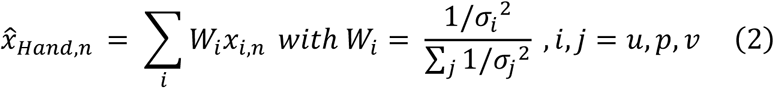

Zhang et al. (2024) found that visual uncertainty (*σ_v_*) linearly scales with error clamp perturbation size, thus the visual uncertainty cue parameter (*σ_v_* =18.706) was determined using experimentally derived slope (*b* = 1.853), and intercept (*a* = 0.309) values from Z. Zhang et al. (2024).

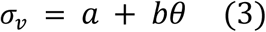

In the model, there were four parameters of interests, namely retention factor (A), error sensitivity (B), sensory prediction noise (*σ_u_*), and proprioceptive noise (*σ_p_*) that were fit with our empirical data. Model fitting was performed in MATLAB using the built-in *fmincon* function, which minimized the root mean square errors (RMSE). To obtain parameter estimates, model fitting was applied using 10,000 bootstrapped samples of our data where the bias-corrected accelerated confidence intervals (CI, 95%) was calculated (Kim, Kitchen, et al., 2025).

Initial conditions for the four free parameters *A, B, σ_u_*, and *σ_p_* were set to 0.95, 0.20, 100, and 25, respectively, with bounds of [0,1] for A and B, and [0, 200] for *σ_u_*, and *σ_p_*. Between group differences in bootstrapped free parameters were determined by examining whether the 95% CIs overlapped or not across AWS and AWNS.

## Results

### Movement duration

We found no statistical difference in movement duration in AWS (*M* = 286 ms, *SD* = 43.410) and AWNS (*M* = 290 ms, *SD* = 48.782), *t*(37.494) = 0.246, *p* = 0.807, *d* = 0.080. The average movement duration in both groups were consistent with prior studies (e.g., Lantagne et al., 2021).

### Adaptation to clamped feedback

Both AWS and AWNS moved their hands more in clockwise direction (relative to the target) in response to clamped feedback (see Fig. 2A). The AWNS control group demonstrated reach adaptation consistent with previously reported levels of approximately 20° of total adaptation under a comparable (i.e., 8° vs 7.5°) clamped feedback paradigm (Morehead et al. 2017).

The linear mixed model for early adaptation revealed a significant fixed effect of group, *F(*1, 40) = 6.618, *p =* 0.014, *ω*^2^ = 0.120, demonstrating that AWS adapted less than AWNS (Fig. 2B, left). There was also significant fixed effects of target, *F*(7, 1880) = 23.700, *p* < 0.001, *ω*^2^ = 0.080, and group × target, *F*(7, 1880) = 2.150, *p =* 0.036, *ω*^2^ = 0.004. Post-hoc pairwise contrasts of the group × target interaction revealed that group differences were found in two targets; 0° direction, *t*(146) = 4.411, *p <* 0.001, *d* = 0.738, and the 45° direction, *t*(146) = 2.520, *p* = 0.013, *d* = 0.457.

AWS also showed reduced adaptation compared to AWNS in late adaptation, *F*(1, 40) = 4.358, *p* = 0.043, *ω*^2^ = 0.070 (Fig. 2B, right). For late adaptation, there were also a significant fixed effect of target, *F*(7, 1880) = 103.347, *p* < 0.001, *ω*^2^ = 0.280 and a significant fixed effect of group × target, *F*(7, 1880) = 5.482, *p <* 0.001, *ω*^2^ = 0.020. Post-hoc pairwise contrasts also revealed that AWS adapted less than AWNS in the 0° direction, *t*(58.6) = 3.804, *p* < 0.001, *d* = 0.827, 45° target direction, *t*(58.6) = 2.620, *p =* 0.011, *d* = 0.714, and 90° direction, *t*(58.6) = 2.622, *p* = 0.027, *d* = 1.201. The fixed effect of targets in both early and late adaptation revealed that both groups generally adapted more in the 0°,45°,180°, and 225° directions (see Supplemental Table S1 and S2 for more details)

### Computational modeling

We fitted adaptation data from AWS and AWNS to the PEA model (Fig. 3A). Average parameter values across our bootstrapped sample fitting for memory retention (A), error sensitivity (B), proprioceptive noise (*σ_p_*), and prediction noise (*σ_u_*) were A = 0.954, B = 0.216, *σ_p_* = 105.709, and *σ_u_* = 160.286 for AWS, and A = 0.960, B = 0.291, *σ_p_* = 102.081, and *σ_u_* = 191.680 for AWNS. 95% bias-corrected accelerated percentile confidence intervals (BCa, *k* = 10,000) were computed for free parameters A, B, *σ_p_*, and *σ_u_* (Fig. 3B).

**Fig. 3.**
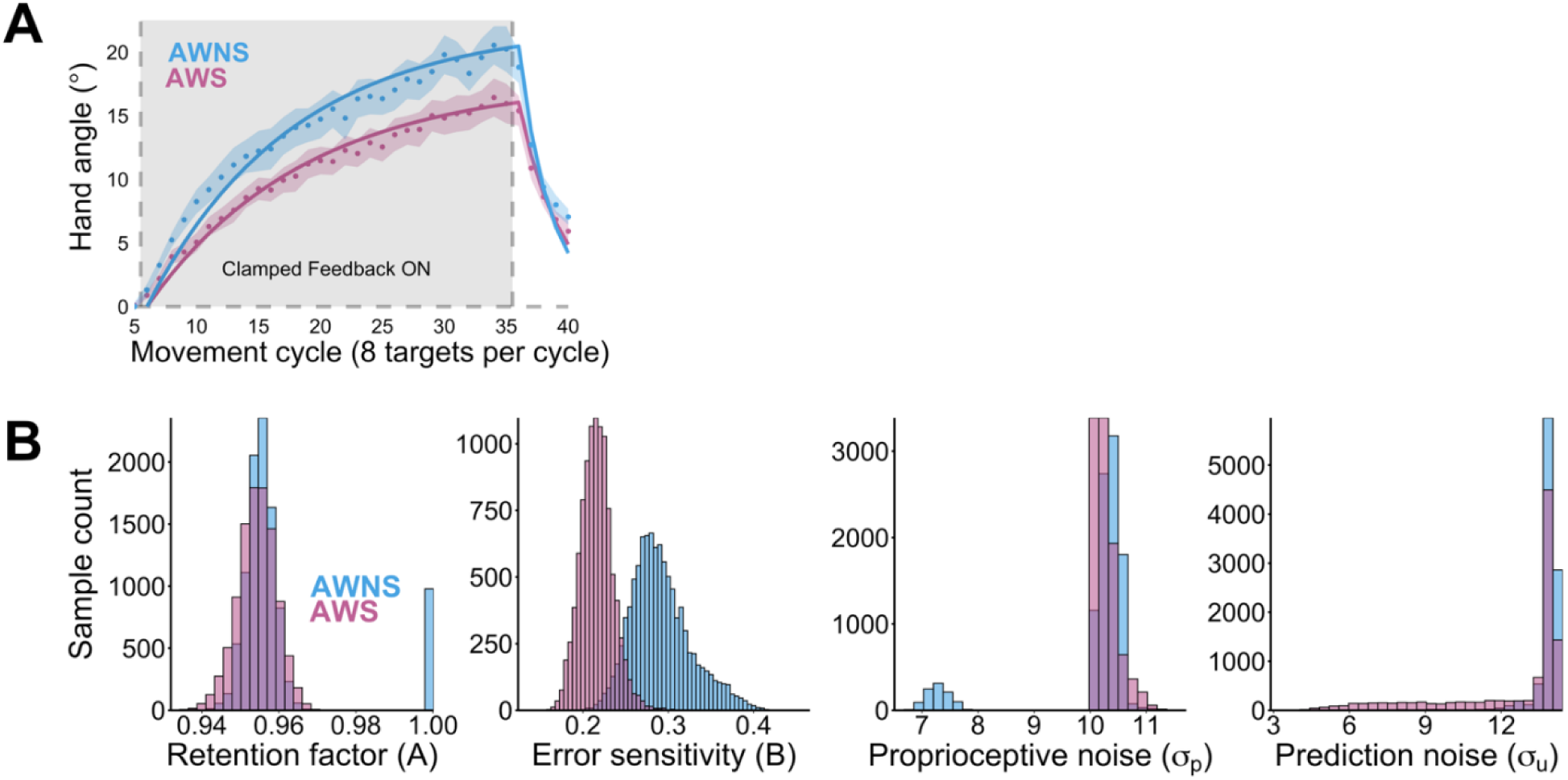
Computational modeling. **(A)** Group averaged adaptation (dots) were fit with the Perceptual Error Attribution (PEA, Zhang et al., 2024) model simulation, generated by average parameter values from 10,000 bootstrapped samples. **(B)** Parameter distributions in the bootstrapped samples showed that retention factor (A), proprioceptive noise (*σ_p_*), and prediction noise (*σ_u_*) had substantial overlap across the two groups, but the error sensitivity (B) had only a slight marginal overlap (< 0.001).

We found that the confidence intervals (CIs) for memory retention (A), proprioceptive noise (*σ_p_*), and prediction noise (*σ_u_*) overlapped substantially across groups (see CIs for the rest of the parameters in Supplemental Table S3), but the error sensitivity parameter (B) showed a marginal (< 0.001) overlap between AWS and AWNS, [0.170, 0.241] vs [0.241, 0.386] (Fig 3.B). To validate our results, we repeated this fitting process for 10 iterations. The CIs marginally overlapped in 8 iterations, but they did not overlap in the remaining 2 iterations, indicating that the trend towards a group difference was consistent across iterations.

## Discussion

The scope of this study was to examine implicit visuomotor adaptation in AWS using a task-irrelevant clamped feedback visuomotor adaptation paradigm which prevents task-based error signals and isolates implicit learning. We found that AWS demonstrated a reduced extent of implicit adaptation compared to AWNS in both early and late adaptation. Our findings suggest that the previously reported slower rate of VMR adaptation in stuttering individuals (Kim et al., 2020) may have been at least partially due to reduced implicit learning difficulties.

To our knowledge, this is the first evidence suggesting that there may be implicit *visuomotor* learning difficulties in people who stutter. Interestingly, it has been reported that AWS also learned less compared to AWNS in *speech* sensorimotor adaptation, an entirely implicit learning task (Keough et al., 2013; Kim & Max, 2021; Lametti et al., 2020). In addition, young children who stutter also showed substantial difficulties in speech adaptation (Kim et al., 2020; but also see Daliri et al., 2018 for a null finding with a different perturbation), suggesting that implicit adaptation difficulties may arise from core sensorimotor deficits associated with stuttering. Our findings build upon this notion by demonstrating that implicit sensorimotor learning difficulties in stuttering individuals may also generalize across multiple effector systems.

A possible explanation for limited implicit adaptation across multiple effector systems could be that there exist general disruptions in their internal forward model (Max et al., 2004; and see also a similar idea in Hickok et al., 2011). Based on this idea, inaccurate sensory predictions generated by the forward model in stuttering individuals may lead to inefficient/incorrect sensory prediction errors. Given that such sensory prediction errors, which reflect discrepancies between forward model predicted sensory outcome and actual sensory input, are known to drive implicit adaptation (Mazzoni & Krakauer, 2006; Taylor et al., 2014), inefficient/incorrect error signals may lead to reduced implicit adaptation in people who stutter. Consistent with this view in the context of speech production, it has been reported that speaking-induced suppression during speech production—a measure of sensory (auditory) prediction errors (Kim, Hinkley, et al., 2025)—is reported to be reduced (Beal et al., 2010) and delayed (Beal et al., 2011) in people who stutter.

Another possible interpretation of our findings is that, perhaps in combination with the disrupted forward model, there is an inefficient updating of control policy (i.e., motor system generating motor commands fine-tuned using error) across multiple effector systems which contributes to limited implicit adaptation. This view is supported by evidence that implicit adaptation may be driven by control policy updates (i.e., updating of feedforward control) rather than forward model updates alone (Hadjiosif et al., 2021; Kim et al., 2023). In line with this view, in the context of speech production, recent reports have highlighted inefficient feedforward control in stuttering individuals. For example, pre-speech auditory modulation, which may reflect inefficient feedforward control (Li et al., 2024), is reduced in stuttering individuals (Daliri & Max, 2015a, 2015b, 2018; Max & Daliri, 2019). In addition, AWS also show more trial-to-trial variability in the early parts of speech movement, measured by initial vowel formant variability, which suggests inefficient feedforward processes (Daliri et al., 2025), consistent with the theoretical perspective (Max et al., 2004).

Our findings also point to a potential role of atypical cerebellar processes in AWS. It has been demonstrated in previous studies that patients with cerebellar degeneration broadly showed impaired visuomotor adaptation (Schlerf et al., 2013), and more specifically attenuated implicit adaptation (Morehead et al., 2017). This interpretation is consistent with converging neuroimaging evidence suggesting that people who stutter have atypical cerebellar activities (Chang et al., 2011; De Nil et al., 2001; Watkins et al., 2008) and disrupted functional connectivity (Chang & Zhu, 2013; Lu et al., 2009). In addition, a recent study identified four subtypes of stuttering using MRI scans in children who stutter and reported a wide range of differences in the cerebellum in children who stutter (Nanda et al., 2026). There were atypical cerebellar volumes in all four subtypes, varying in the direction (i.e., increase vs. decrease) and location depending on the subtypes. Together, these findings add support to the overall notion that atypical cerebellar anatomies and functions may play a key role in the implicit sensorimotor learning differences observed in AWS.

### Computational modeling

To identify specific processes underlying adaptation difficulties, we fit group averaged adaptation across targets to the PEA model which assumes that implicit sensorimotor adaptation is driven by a combination of proprioceptive, sensory prediction, and visual error cues in a Bayesian manner (Z. Zhang et al., 2024). Within our 10,000 bootstrapped samples, we found substantial CI overlap between the two groups (AWS vs. AWNS) in proprioceptive, sensory prediction, and memory retention rate parameters, but interestingly only a marginal overlap (< 0.001) in error sensitivity, potentially suggesting that AWS may have a reduced sensitivity to errors (see Fig. 3B). Such reduced error sensitivity is consistent with the potential interpretations with an inaccurate forward model and/or poor control policy updates in stuttering, but some cautions should be taken before interpreting this modeling results. First, given the marginal overlap in CIs for error sensitivity, future studies should rigorously test each parameter independently to obtain more precise values and further validate our modeling results. In addition, even if reduced error sensitivity is later identified as a source of limited adaptation in stuttering individuals, it should be noted that error sensitivity is typically approximated from a subsequent movement following the experience of errors, an outcome that confounds multiple processes such as error detection, error evaluation, and incorporation of the errors in motor planning for the subsequent movements. As a result, more investigations are necessary to understand which process(es) lead to reduced error sensitivity.

### Differences across movement directions

Beyond the primary findings of group differences, we also observed significant fixed effects in target and group × target interactions. Across early and late adaptation, we found that between-group adaptation differences were dependent on reach direction. In our post-hoc analyses, the group difference occurred in both the 0° and 45° reach directions. Interestingly, the adjacent 90° direction also had statistically significant group differences but only for late adaptation. Given that these three directions are adjacent to each other, it is possible that updating motor commands for this cluster of directions may be more challenging for AWS. For example, it is possible that targets in the 0°, 45°, and 90° directions require more complex motor plan updates given that they corresponded to reaching directions associated with moderate-to-high inertia (Gordon et al., 1994). However, we did not observe group differences in the other directions with the same amount of inertia (e.g., 225°), suggesting that inertia alone may not explain our findings. Alternatively, three directions showing significant group differences may reflect a random spurious effect, given that, on average, AWS showed less adaptation in all eight movement directions compared to AWNS (Fig. 2 and Tables S2, S3).

### Future directions

A few limitations of our study should be acknowledged. First, estimations of explicit and implicit learning highly depend on methods (Hadjiosif et al., 2021; Hadjiosif & Krakauer, 2021), suggesting that our findings should be validated in different tasks involving different instructions and contexts. Second, because the current study did not assess any explicit components, it cannot be argued, based on our findings, that limited visuomotor adaptation found in AWS reflects *entirely* implicit learning difficulties. In other words, it remains possible that visuomotor adaptation difficulties in AWS found in Kim et al. (2020) may partially reflect poor explicit learning. Future studies are warranted to measure explicit learning in AWS using both behavioral task (Taylor et al., 2014) and computational modeling (X. Zhang et al., 2026).

In addition, we previously found that both children who stutter and children who do not stutter (under 10 years of age) adapted a relatively small amount to the VMR and hypothesized that the sensorimotor system in children may not be mature enough to show adult-like adaptation (Kim et al., 2020). It is possible that the clamped feedback adaptation paradigm used in the current study may be easier for younger children’s sensorimotor system. Future studies are warranted to examine whether implicit visuomotor adaptation difficulties are also present in children who stutter given that stuttering is a developmental disorder.

## Conclusion

The current study demonstrates that stuttering individuals show implicit visuomotor adaptation difficulties. Computational modeling using the PEA model also identified error sensitivity as a potential factor that may contribute to the limited implicit adaptation in people who stutter. Overall, our findings provide support for the idea that stuttering may be associated with unstable/inefficient internal models that affect multiple effector systems. Future work should validate and expand upon our findings to determine specific processes that contribute to limited implicit adaptation (and perhaps reduced error sensitivity) in stuttering.

## Data Availability

Our dataset for this study has been uploaded to an open-source repository that can be accessed at https://osf.io/c5s7w/

## Acknowledgements

We would like to thank Anabelle Ross and Barbara Brown for their help with stuttering severity index. We would also like to thank the National Stuttering Association West Lafayette chapter for their support for participant recruitment.

## Supplemental Information

**Table S1.** Estimated marginal means (EMMs) for early adaptation for each target and group.

| Target | Group | Estimate | Lower 95% CI | Upper 95% CI |
| --- | --- | --- | --- | --- |
| **Target 1 (45°)** | AWNS | 7.64 | 6.20 | 9.08 |
|  | AWS | 5.04 | 3.60 | 6.48 |
| **Target 2 (90°)** | AWNS | 3.76 | 2.32 | 5.20 |
|  | AWS | 2.14 | 0.70 | 3.58 |
| **Target 3 (135°)** | AWNS | 3.68 | 2.24 | 5.12 |
|  | AWS | 1.82 | 0.38 | 3.27 |
| **Target 4 (180°)** | AWNS | 6.49 | 5.05 | 7.93 |
|  | AWS | 5.53 | 4.09 | 6.97 |
| **Target 5 (225°)** | AWNS | 6.44 | 5.00 | 7.88 |
|  | AWS | 4.95 | 3.51 | 6.39 |
| **Target 6 (270°)** | AWNS | 4.10 | 2.66 | 5.54 |
|  | AWS | 3.26 | 1.82 | 4.70 |
| **Target 7 (315°)** | AWNS | 3.87 | 2.43 | 5.31 |
|  | AWS | 2.24 | 0.80 | 3.68 |
| **Target 8 (0°)** | AWNS | 9.44 | 8.00 | 10.89 |
|  | AWS | 5.20 | 3.76 | 6.64 |

*Note. SE = 0.73, df = 146 for all early comparisons (Kenward-Roger). AWNS = adults with no stutter;*
*AWS = adults who stutter. 95% confidence intervals shown.*

**Table S2.** Estimated marginal means (EMMs) for late adaptation for each target and group.

| Target | Group | Estimate | Lower 95% CI | Upper 95% CI |
| --- | --- | --- | --- | --- |
| **Target 1 (45°)** | AWNS | 25.6 | 22.47 | 28.6 |
|  | AWS | 19.9 | 16.77 | 22.9 |
| **Target 2 (90°)** | AWNS | 16.1 | 12.97 | 19.1 |
|  | AWS | 11.1 | 8.04 | 14.2 |
| **Target 3 (135°)** | AWNS | 13.5 | 10.41 | 16.6 |
|  | AWS | 11.2 | 8.13 | 14.3 |
| **Target 4 (180°)** | AWNS | 21.2 | 18.11 | 24.3 |
|  | AWS | 18.2 | 15.11 | 21.3 |
| **Target 5 (225°)** | AWNS | 24.2 | 21.09 | 27.2 |
|  | AWS | 20.8 | 17.75 | 23.9 |
| **Target 6 (270°)** | AWNS | 16.7 | 13.66 | 19.8 |
|  | AWS | 14.3 | 11.24 | 17.4 |
| **Target 7 (315°)** | AWNS | 14.1 | 11.03 | 17.2 |
|  | AWS | 11.3 | 8.25 | 14.4 |
| **Target 8 (0°)** | AWNS | 25.9 | 22.80 | 29.0 |
|  | AWS | 17.6 | 14.52 | 20.7 |

*Note. SE = 1.54, df = 58.6 for all late comparisons (Kenward-Roger). AWNS =adults with no stutter;*
*AWS adults who stutter. 95% confidence intervals shown.*

**Table S3.**
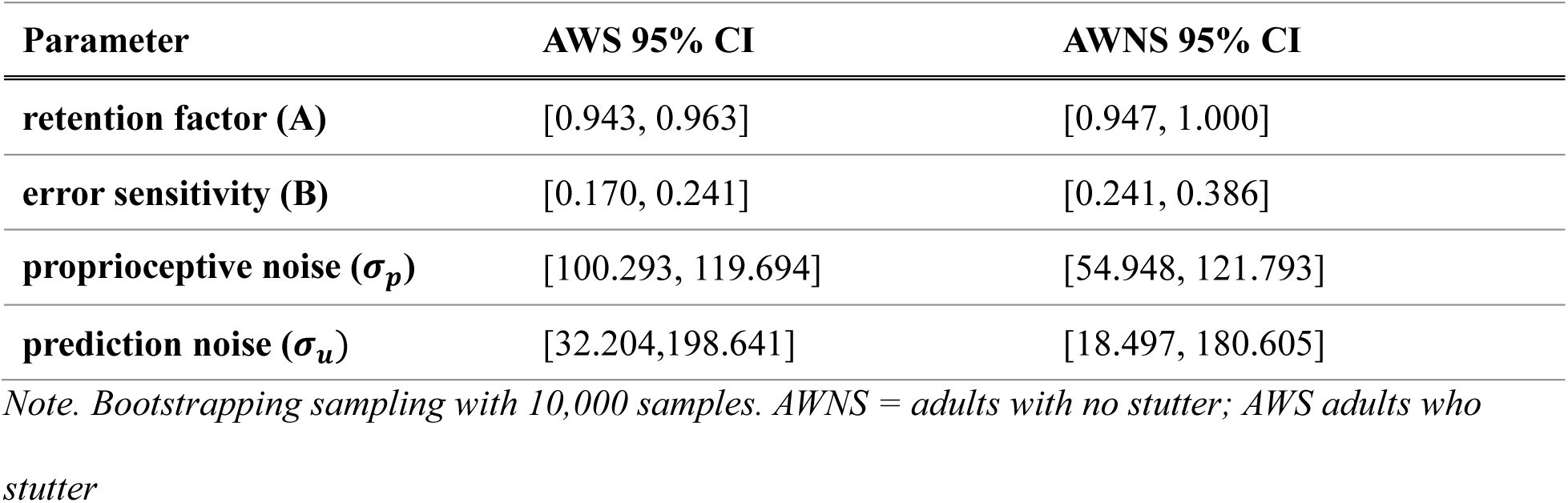
Modeling results by group.

| **Parameter** | **AWS 95% CI** | **AWNS 95% CI** |
| --- | --- | --- |
| **retention factor (A)** | [0.943, 0.963] | [0.947, 1.000] |
| **error sensitivity (B)** | [0.170, 0.241] | [0.241, 0.386] |
| **proprioceptive noise ($\sigma_p$)** | [100.293, 119.694] | [54.948, 121.793] |
| **prediction noise ($\sigma_u$)** | [32.204, 198.641] | [18.497, 180.605] |

*Note. Bootstrapping sampling with 10,000 samples. AWNS = adults with no stutter; AWS adults who stutter*

